# Detection of genes with differential expression dispersion unravels the role of autophagy in cancer progression

**DOI:** 10.1101/2022.07.01.498392

**Authors:** Christophe Le Priol, Chloé-Agathe Azencott, Xavier Gidrol

**Affiliations:** Univ. Grenoble Alpes, INSERM, CEA-IRIG, Biomics, Grenoble, France; Center for Computational Biology, Mines ParisTech, PSL Research University, Paris, France; Institut Curie, Paris, France; INSERM U900, Paris, France

## Abstract

The majority of gene expression studies focus on the search for genes whose mean expression is different between two or more populations of samples in the so-called “differential expression analysis” approach. However, a difference in variance in gene expression may also be biologically and physiologically relevant. In the classical statistical model used to analyze RNA-sequencing (RNA-seq) data, the dispersion, which defines the variance, is only considered as a parameter to be estimated prior to identifying a difference in mean expression between conditions of interest. Here, we propose to evaluate two recent methods, MDSeq and DiPhiSeq, which detect differences in both the mean and dispersion in RNA-seq data. We thoroughly investigated the performance of these methods on simulated datasets and characterized parameter settings to reliably detect genes with a differential expression dispersion. We applied both methods to The Cancer Genome Atlas datasets. Interestingly, among the genes with an increased expression dispersion in tumors and without a change in mean expression, we identified some key cellular functions, most of which were related to catabolism and were overrepresented in most of the analyzed cancers. In particular, our results highlight autophagy, whose role in cancerogenesis is context-dependent, illustrating the potential of the differential dispersion approach to gain new insights into biological processes.

**Author summary:** Gene expression is the process by which genetic information is translated into functional molecules. Transcription is the first step of this process, consisting of synthesizing messenger RNAs. During recent decades, genome-wide transcriptional profiling technologies have made it possible to assess the expression levels of thousands of genes in parallel in a variety of biological contexts. In statistical analyses, the expression of a gene is estimated by counting sequencing reads over a set of samples and is defined by two dimensions: mean and variance. The overwhelming majority of gene expression studies focus on identifying genes whose mean expression significantly changes when comparing samples of different conditions of interest to gain knowledge of biological processes. In this classical approach, the variance is usually considered only as a noise parameter to be estimated before assessing the mean expression. However, finely estimating the variance of expression may be biologically relevant since a modification of this parameter may reflect a change in gene expression regulation. Here, we propose to evaluate the performance of statistical methods that identify such differentially variant genes. We highlighted the potential of this approach by analyzing cancer datasets, thus identifying key cellular functions in tumor progression.

## Introduction

### Variability in gene expression in cancer

Genome-wide transcriptional profiling technologies have made it possible to assess the level of expression of thousands of genes in parallel in a variety of biological contexts [1]. Cells or organs are commonly characterized by the mean expression of some key genes [2]. As a consequence, phenotypes are defined to be driven by a change in the mean expression of some genes between sets of samples that represent conditions of biological interest, *e.g*. diseased and healthy status [3]. Several methods have thus been developed to identify these genes, called “differentially expressed” (DE) genes. This has led to numerous insights into a variety of biological processes [4, 5]. Differentially expressed genes may also serve as biomarkers [6]. In this type of analysis, the variability is often reduced to “noise” that one must remove. Consequently, variability is considered to be a parameter that must be estimated prior to searching for a difference in mean expression. However, in the same manner that the level of expression of a gene has biological meaning, the variability of its expression is another trait of its biological function [7, 8]. For example, low gene expression variability defines housekeeping genes [9, 10] and is a desirable property when identifying reliable biomarkers [11].

The fluctuations in gene expression may indeed be driven by a variety of intrinsic sources, *e.g*. the stochastic nature of gene transcription [12], the cell cycle [13], stochastic regulation [14], chromatin modification [15] or mRNA degradation [16], as well as extrinsic causes, which refer to all environmental perturbations [17, 18]. In cancer, the overall increase in gene expression variability [19] is a way for tumors to resist therapy [20, 21]. In addition, it may reveal a lack of precision in gene expression, which tends to be highly controlled in healthy conditions [22, 23]. For these reasons, variability is a relevant trait in gene expression to gain better knowledge of cancer development.

### Statistical analysis of gene expression variability

The terms “variability” and “variation” are often used to describe how much the expression of a gene fluctuates when comparing different samples. These terms may be confusing when analyzing samples from different biological conditions, since they are commonly used to refer to a change of mean expression between conditions. In addition, they are not statistical terms and should therefore be replaced by the metric used to estimate the variability in the analyzed data. A myriad of measures may be used to estimate gene expression variability, *e.g*. the variance, the standard deviation, the coefficient of variation (CV), the median absolute deviation, the expression variability [10], the Shannon entropy [24] or the expression change [22].

Genes having a difference of variance in expression between biological conditions of interest are called “differentially variant” (DV) genes and are identified using basic statistical approaches: F-test to compare variances [8, 25], Wilcoxon rank-sum test to compare CVs [26, 27], differences of entropy tests [24] or comparison of CV distributions to random distributions using Wilcoxon’s signed rank test [28]. A few studies have focused on analyzing gene expression variability and identified genes with differential variance in different biological contexts: cancer progression [8, 25], neurologic diseases such as Parkinson’s disease and schizophrenia [26, 29] or between cell populations in development [27]. Most of these studies used microarrays and log-transformed the expression data prior to measuring gene expression variability. This transformation affects the mean-variance relationship [30] and therefore appears to be suboptimal for estimating gene expression variability.

High-throughput sequencing of the transcriptome (RNA-seq) has become the gold-standard technology to estimate genome-wise gene expression [31]. Contrary to microarray data, RNA-seq count data are integer values, which makes log-transformation, usually performed with microarray data, not appropriate for this type of data [32]. Therefore, dedicated methods based on discrete probability distributions were developed to analyze these data [33]. The negative binomial (NB) distribution has become the ubiquitous distribution to model RNA-seq read count data by providing the best fit for the extra-variance commonly observed in datasets composed of biological replicates [34]. In this model, the random variable describing the count of reads mapped to gene *i* in sample *j* is denoted as *Y*_*ij*_ ∼ *𝒩 ℬ*(*μ*_*ij*_, *ϕ*_*i*_), where *μ*_*ij*_ is the expected value and *ϕ*_*i*_ is the dispersion parameter. The variance is given by 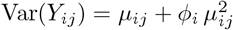. Analyzing the variance independently with respect to the mean expression can therefore be achieved by analyzing the dispersion parameter *ϕ*_*i*_.

In the classical RNA-seq data analysis workflow, differential expression detection methods based on the NB distribution consider the dispersion as a noise parameter to be estimated prior to identifying a difference of mean expression [35]. The generally low sample sizes of RNA-seq datasets at the time when the first versions of these methods were published made per-gene dispersion estimation unreliable. In addition, the very high number of genes made estimation difficult. Thus, Robinson *et al*. proposed an accurate shared estimator based on the expression of sets of genes across all samples, independent of biological condition [34]. Per-gene estimators were then shrunk towards this shared estimator using different levels of shrinkage [36–39]. Aggregating all the samples that compose the dataset implies that no difference of dispersion in the expression of genes between the conditions of interest can be modeled, which is not biologically realistic.

Recently, two new methods based on the NB distribution, MDSeq [40] and DiPhiSeq [41], have been introduced to identify differences in both mean and dispersion in RNA-seq data within the same statistical framework. MDSeq extends the use of a generalized linear model (GLM) to identify both mean and dispersion differences by reparameterizing the NB distribution with a linear mean-variance relationship: Var(*Y*_*ij*_) = *ϕ*_*ij*_ *μ*_*ij*_. Since the NB distribution with a varying dispersion parameter does not belong to the exponential family, the usual closed-form estimates for the GLM parameters cannot be used. Instead, the minimization of the log-likelihood of the model is formulated as an optimization problem with linear inequality constraints that can be solved using an adaptive barrier algorithm combined with the BFGS algorithm. Wald tests were then performed to identify differential expression mean and dispersion. DiPhiSeq does not implement a GLM but, unlike the classical differential expression methods, estimates the dispersion for each gene and for the two compared conditions. Because of the high sensitivity of the likelihood ratio test to outliers, the authors of DiPhiSeq used robust M-estimators to estimate both the mean and the dispersion in both conditions. In this approach, the Tukey’s biweight function is used as the function to minimize. Differences in the mean and dispersion are finally compared to a standard distribution under the null hypothesis of no difference and p-values for differential expression and differential dispersion are deduced.

### Objectives

The performances of methods identifying differences in mean expression in the so-called “differential expression analysis” using RNA-seq data have been extensively studied [42–45]. The large amount of publicly available RNA-seq data opens new perspectives for researchers in the search for genes whose expression exhibits a difference of dispersion between samples from different conditions. Here, we propose to evaluate the performances of two NB-based methods, MDSeq and DiPhiSeq, to identify differentially dispersed (DD) genes using simulated RNA-seq datasets. In their respective articles, these methods were not compared to each other, but rather to non-NB-based differential variance methods for MDSeq and to NB-based differential expression methods for DiPhiSeq to highlight the biological interest of identifying DD genes. Based on our simulation study results, we reliably applied these methods to The Cancer Genome Atlas datasets and identified DD genes that could not be identified by classical differential expression analysis. We showed that these genes may serve to better understand tumor progression and thus have demonstrated the potential of the differential dispersion approach in RNA-seq studies.

## Results

### DiPhiSeq and MDSeq performance evaluation

#### Differential dispersion detection for genes with unconstrained differences in mean expression

We simulated RNA-seq datasets to evaluate the performances of DiPhiSeq and MDSeq to identify differential dispersion between two sets of samples of equal size that represent two conditions of interest. Differences in the mean and dispersion between the two sets of samples were introduced and defined for DE and DD genes, respectively (see the Methods section for more details). DiPhiSeq achieves a better overall performance for differential dispersion detection than MDSeq regardless of the number of samples per condition (red boxplots in Fig 1).

**Fig 1.**
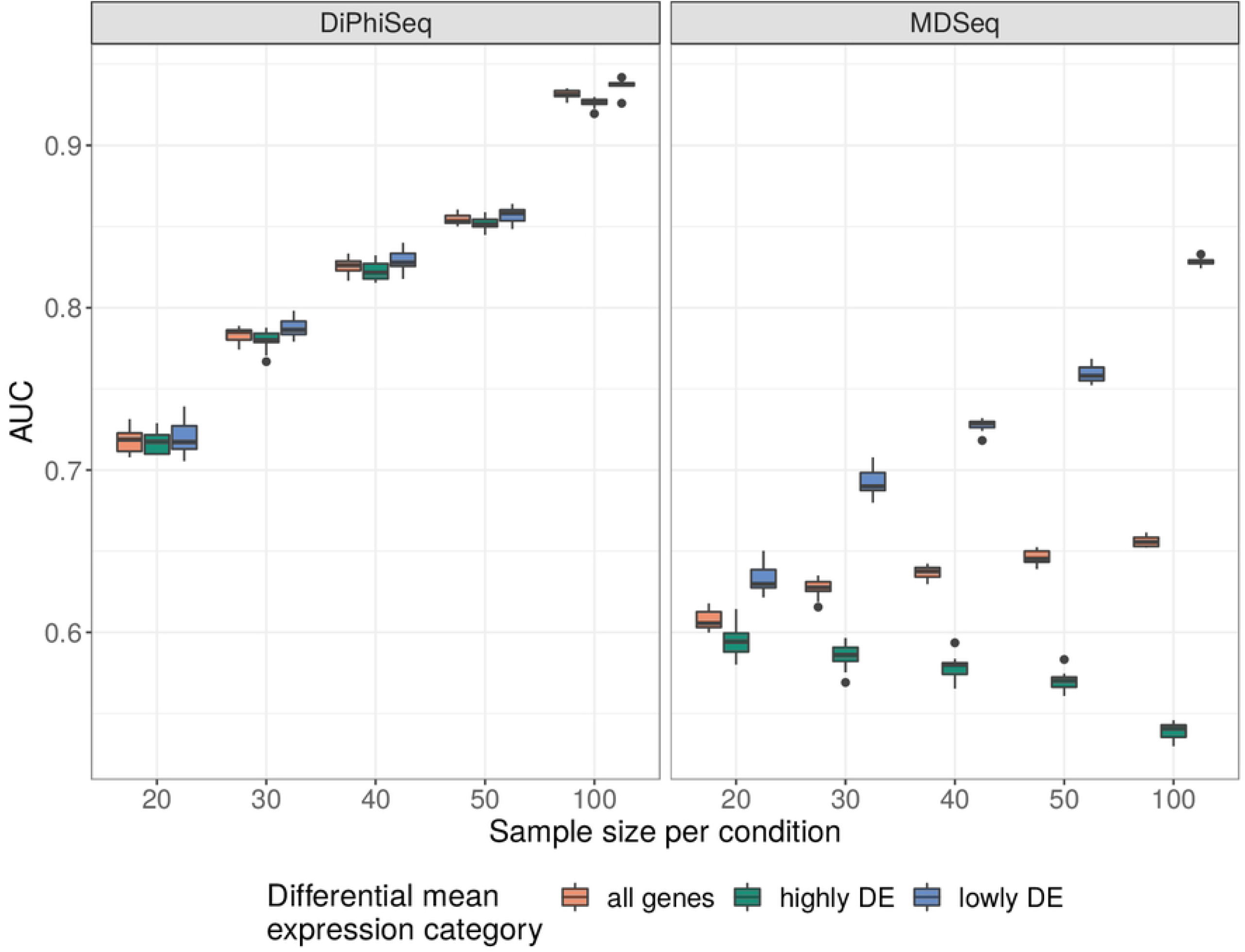
DiPhiSeq and MDSeq ability to identify differentially dispersed genes. DiPhiSeq has a better ability to identify differentially dispersed genes, as measured by the area under the ROC curve (AUC), than MDSeq, whose performance declines when the genes are also differentially expressed.

This noticeable difference may be explained by the difference in false discovery rate (FDR) controlling procedures used by the two methods: the Benjamini-Hochberg procedure for DiPhiSeq and the Benjamini-Yekutieli procedure for MDSeq, as recommended by the authors of the two methods. The Benjamini-Yekutieli procedure is more conservative than the former [46] and thus may explain the lower area under the ROC curve (AUC) and sensitivity values obtained with MDSeq. We note, however, that in our evaluation (see S2 Fig), the Benjamini-Hochberg procedure was not sufficient to control for FDR.

As expected, increasing the number of samples available per condition increases the ability to detect differential dispersion. Nevertheless, these sample sizes are much larger than those usually required to achieve similar performances in classical differential expression analysis [42–44]. For example, 40 samples per condition are required for DiPhiSeq to obtain an AUC higher than 0.8, and sets of 50 samples are required for MDSeq to obtain an AUC close to this value among lowly DE genes, while only 5 samples may suffice to identify differences in mean expression with this performance [44].

The other main result of our simulation study is that a fold-change in the mean sharply reduces the performance of MDSeq for differential dispersion detection. By contrast, DiPhiSeq is not sensitive to the presence of a difference in mean expression between the two compared sets of samples. The AUC obtained with MDSeq can indeed be as much as 20% lower when the genes are also highly DE (green vs. blue boxplots in Fig 1). The application of MDSeq to identify differential dispersion must therefore be restricted to non- or lowly DE genes.

#### Differential dispersion detection for lowly DE genes

The maximum difference in means of gene expression according to the number of samples in the two compared conditions while maintaining the reliability of the differential dispersion detection with MDSeq must therefore be identified. Given the results in Fig 1, this number is expected to depend on the number of samples. Fig 2 shows the performances of MDSeq and DiPheSeq on simulated datasets stratified by the maximum tolerated mean expression fold-change value and the number of samples per condition.

**Fig 2.**
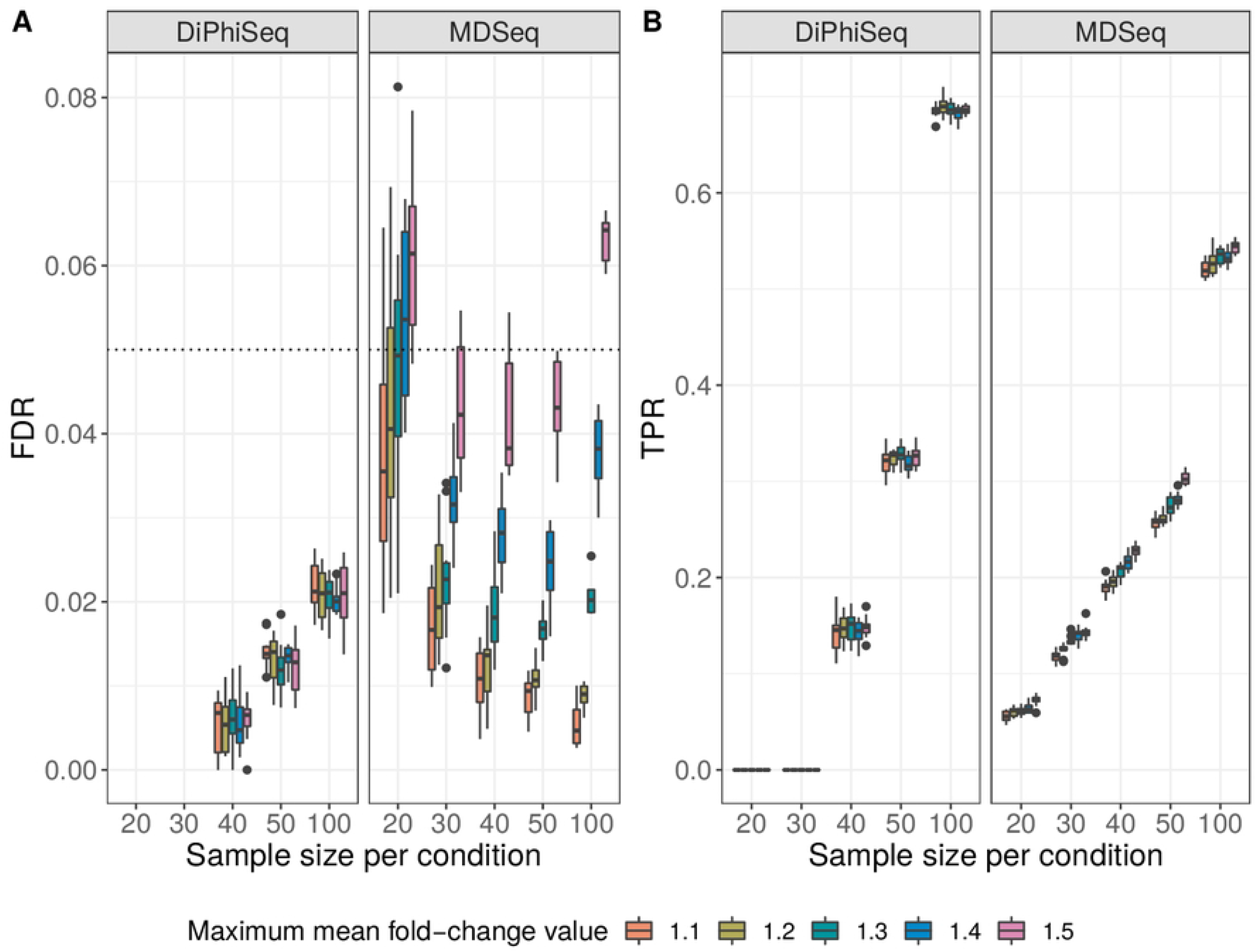
DiPhiSeq and MDSeq ability to detect differential dispersion for lowly differentially expressed genes. (A) False discovery rate (FDR). (B) True positive rate (TPR). The performances of DiPhiSeq and MDSeq for differential dispersion detection in gene expression data were assessed using simulated datasets composed of lowly differentially expressed genes between two sample populations of equal size.

For MDSeq, increasing the maximum tolerated mean fold-change value increases the FDR for the detection of differential dispersion. However, the FDR remained under 0.05 for datasets composed of 30 to 50 samples with maximum tolerated mean fold-changes up to 1.5 (Fig 2A). When only 20 samples are available, the maximum tolerated mean fold-change value must be at most 1.3 to keep the FDR below 0.05.

As already shown in Fig 1, the performance of DiPhiSeq is not affected by the maximum tolerated mean fold-change value (Fig 2A and B). However, DiPhiSeq is less sensitive than MDSeq for low sample sizes (under 40 samples). DiPhiSeq is indeed unable to detect any DD genes with fewer than 30 samples (Fig 2B). In contrast, regarding larger sample sizes, *i.e*. populations of at least 50 samples, DiPhiSeq has a better sensitivity than MDSeq, with an even larger gain in sensitivity as the sample size increases. Overall, DiPhiSeq and MDSeq exhibit close and complementary performances for the differential dispersion detection of lowly DE genes.

This limitation in the application of MDSeq is not prohibitive since the purpose of our approach is to identify genes that would not be detected by the classical differential expression analysis or, at least, that would not appear in the top results of these analyses. Thus, in our approach, lowly DE genes represent the set of genes of primary interest among which to search for differential dispersion in expression. To apply MDSeq, highly DE genes must therefore be filtered out by using a fold-change threshold prior to detecting differential dispersion among the genes that have passed this filter. MDSeq provides the possibility to use specific threshold values to identify both DE and DD genes. We therefore used a range of different mean fold-change threshold values to filter highly DE genes and evaluated MDSeq differential dispersion performance with respect to the genes that passed the filter. The maximum mean fold-change threshold values that enable an increase in the sensitivity of the differential dispersion detection while maintaining the FDR below 0.05 were identified according to several sample sizes (S3 Fig). For example, when comparing two populations of samples of equal size from 30 to 50 samples, a maximum mean fold-change threshold of 1.30 should be used to filter highly DE genes, whereas this maximum threshold should be lowered to 1.15 and 1.25 for populations of 20 and 100 samples, respectively.

The analysis of the true DD genes identified with DiPhiSeq and MDSeq among lowly DE genes when comparing populations of 50 samples revealed that approximately half (54%) of true positive results were identified by both methods, which represents 67.9% and 72.5% of the overall true positives identified by DiPhiSeq or MDSeq, respectively (Fig 3A).

**Fig 3.**
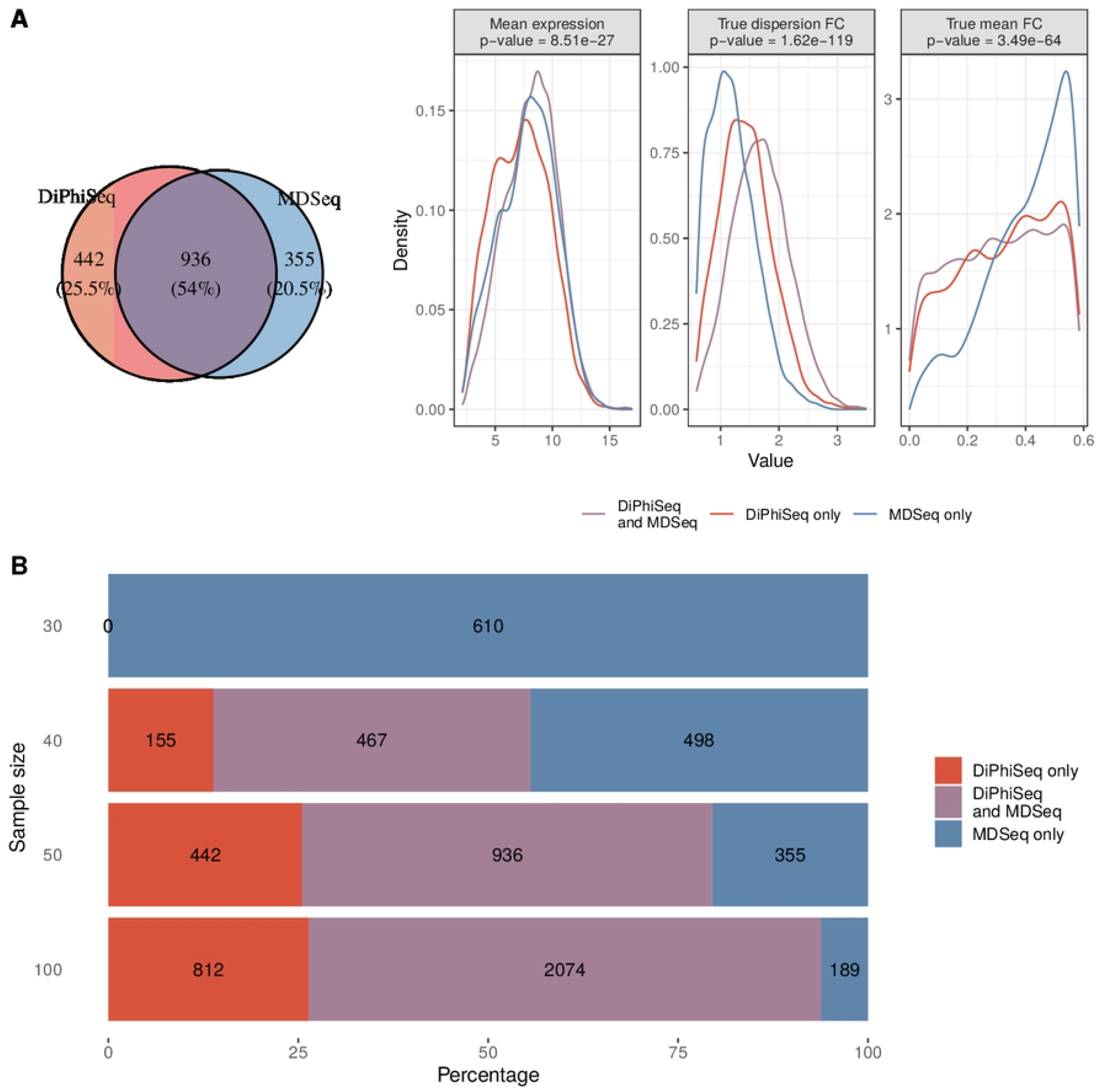
True DD genes identified by either DiPhiSeq, MDSeq, or both among lowly differentially expressed genes. (A) Numbers of true differentially dispersed (DD) genes identified by DiPhiSeq, MDSeq, or both (Venn diagram), as well as histograms of mean expression and absolute values of true dispersion and mean log_2_-fold-changes, over 10 replicates of simulated datasets composed of two populations of 50 samples. Two-sample Wilcoxon tests were performed to evaluate whether these statistics were greater for MDSeq-specific DD genes (mean expression and absolute values of true mean log_2_-fold-changes) or for DiPhiSeq-specific DD genes (absolute values of true dispersion log_2_-fold-changes). P-values of these tests are indicated in panel titles. (B): Numbers of true DD genes identified by DiPhiSeq, MDSeq, or both over 10 replicates of simulated datasets of 20 to 100 samples.

Nevertheless, the numbers of true DD genes identified by only DiPhiSeq or MDSeq, 442 and 355 on average for the datasets in Fig 3A, represent substantial gene sets that cannot be neglected. Since the FDR is guaranteed to be lower than 0.05 for both methods according to our simulation study, the DD genes identified by at least one of the two methods should be kept for subsequent analysis, in addition to those identified by both methods, to gain more biological insight. In addition, specific characteristics of each method can be determined from the DD genes that are identified by one and only one of them. DiPhiSeq-specific DD genes have lower mean expression than MDSeq-specific DD genes, revealing a higher sensitivity of DiPhiSeq for differential dispersion detection among lowly expressed genes (Fig 3A). MDSeq detects lower differences in dispersion in comparison with DiPhiSeq and tends to detect DD genes among genes having a mean fold-change close to the tolerated maximum value (Fig 3A). When 50 or fewer samples are available per set, combining the results of the two methods is relevant since the DD genes detected by both methods only represent at most 54% of the overall DD genes (Fig 3B). As the number of samples per condition increases, the proportion of DD genes specifically detected with MDSeq decreases, whereas the proportion of DiPhiSeq-specific DD genes increases. Regarding larger populations composed of 100 samples, most of the DD genes were detected by both methods (67.4%), while the maximum proportion of DD genes detected with DiPhiSeq reached 26.4%.

### Differential dispersion in gene expression in cancer

Having defined the optimal conditions of utilization, we applied DiPhiSeq and MDSeq to The Cancer Genome Atlas (TCGA) datasets [47] to identify DD genes when comparing normal and tumor samples. We used RNA-seq data from patients for whom tumor tissue and adjacent normal tissue samples were available. In agreement with the results of our simulation study, only the datasets with more than 30 samples for both conditions were analyzed, in order to maintain FDR below 0.05 with MDSeq and to ensure sufficient power with DiPhiSeq. We list these datasets in Table 1.

**Table 1.** Numbers of normal and tumor samples for the analyzed TCGA datasets.

| Dataset | Samples |  |
| --- | --- | --- |
|  | normal | tumor |
| TCGA-BRCA | 112 | 117 |
| TCGA-COAD | 41 | 46 |
| TCGA-HNSC | 43 | 43 |
| TCGA-KIRC | 72 | 72 |
| TCGA-KIRP | 31 | 31 |
| TCGA-LIHC | 50 | 50 |
| TCGA-LUAD | 57 | 67 |
| TCGA-LUSC | 49 | 49 |
| TCGA-PRAD | 52 | 54 |
| TCGA-THCA | 58 | 58 |

The samples originate from patients for whom samples of tumor tissues and adjacent normal tissues are available. Only the datasets with at least 30 samples for both conditions are analyzed: BRCA (BReast invasive CArcinoma), COAD (COlon ADenocarcinoma), HNSC (Head and Neck Squamous cell Carcinoma), KIRC (KIdney Renal Clear cell carcinoma), KIRP (KIdney Renal Papillary cell carcinoma), LIHC (LIver Hepatocellular Carcinoma), LUAD: (LUng ADenocarcinoma), LUSC (LUng Squamous cell Carcinoma), PRAD (PRostate ADenocarcinoma), and THCA (THyroid CArcinoma). For some datasets, the numbers of samples from normal and tumor tissues are different because several samples from tumors are available and are integrated in the analysis.

#### Identification of DD genes

A fold-change threshold of 1 was used to filter DE genes and identify DD genes among non-DE genes with DiPhiSeq and MDSeq. Fig 4 shows the number of DE and DD genes identified for each dataset among non-DE genes. The numbers of DD genes detected irrespective of DE status are reported in S4 Fig and S5 Fig.

**Fig 4.**
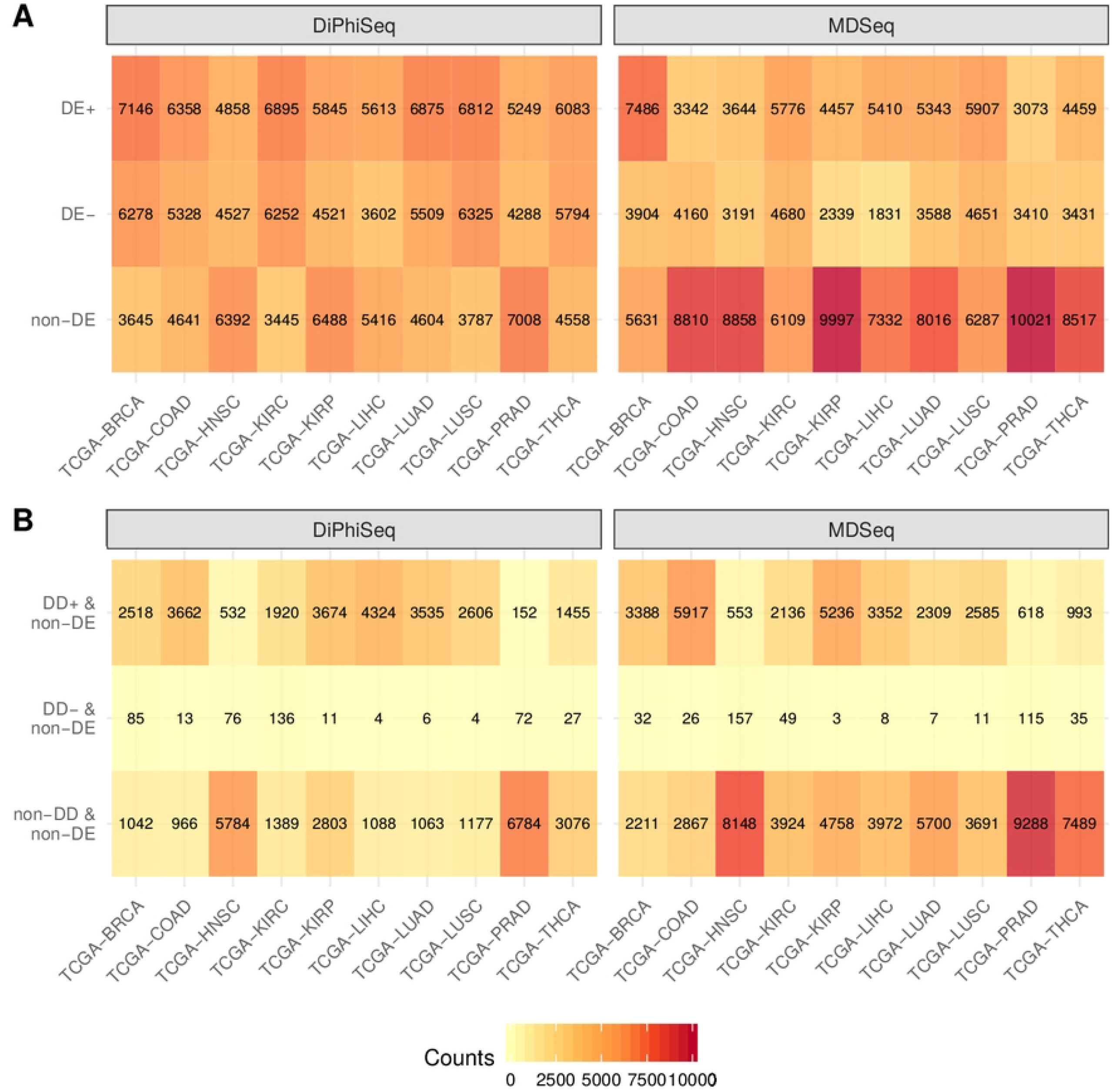
Differentially expressed and differentially dispersed genes according to DiPhiSeq and MDSeq for each TCGA dataset. (A) Number of differentially expressed (DE) genes separated between those upregulated in tumors (DE+) and those downregulated in tumors (DE-), as detected by DiPhiSeq and MDSeq, per TCGA dataset. (B) Number of differentially dispersed (DD) genes among non-DE genes separated between those overdispersed in tumors (DD+) and those underdispersed in tumors (DD-), as detected by DiPhiSeq and MDSeq, per TCGA dataset.

Many more genes are identified as DE by DiPhiSeq than by MDSeq, which dramatically reduces the set of genes of interest among which to search for DD genes. More specifically, there were between 3 445 and 7 008 non-DE genes according to DiPhiSeq and between 5 631 and 10 021 non-DE genes according to MDSeq, depending on the dataset. Nevertheless, there are several thousand genes among which DD genes can be searched for in any dataset. Among non-DE genes, the majority of DD genes are overdispersed in tumors (DD+). Both methods generate consistent results: some cancers are characterized by a high number of DD+ genes (breast, colon, kidney, liver and lung), and others only contain very few DD genes (head and neck, prostate and thyroid).

To use the same gene sets of comparison and because MDSeq cannot identify a differential dispersion in the expression data of highly DE genes while maintaining the FDR below 0.05 according to our simulation study, we compared the DD+ genes identified with DiPhiSeq and MDSeq among the genes that are non-DE according to MDSeq. We observed the same trend as in the simulation study with TCGA datasets: the sets of DD+ genes identified with DiPhiSeq and MDSeq were quite consistent (Fig 5).

**Fig 5.**
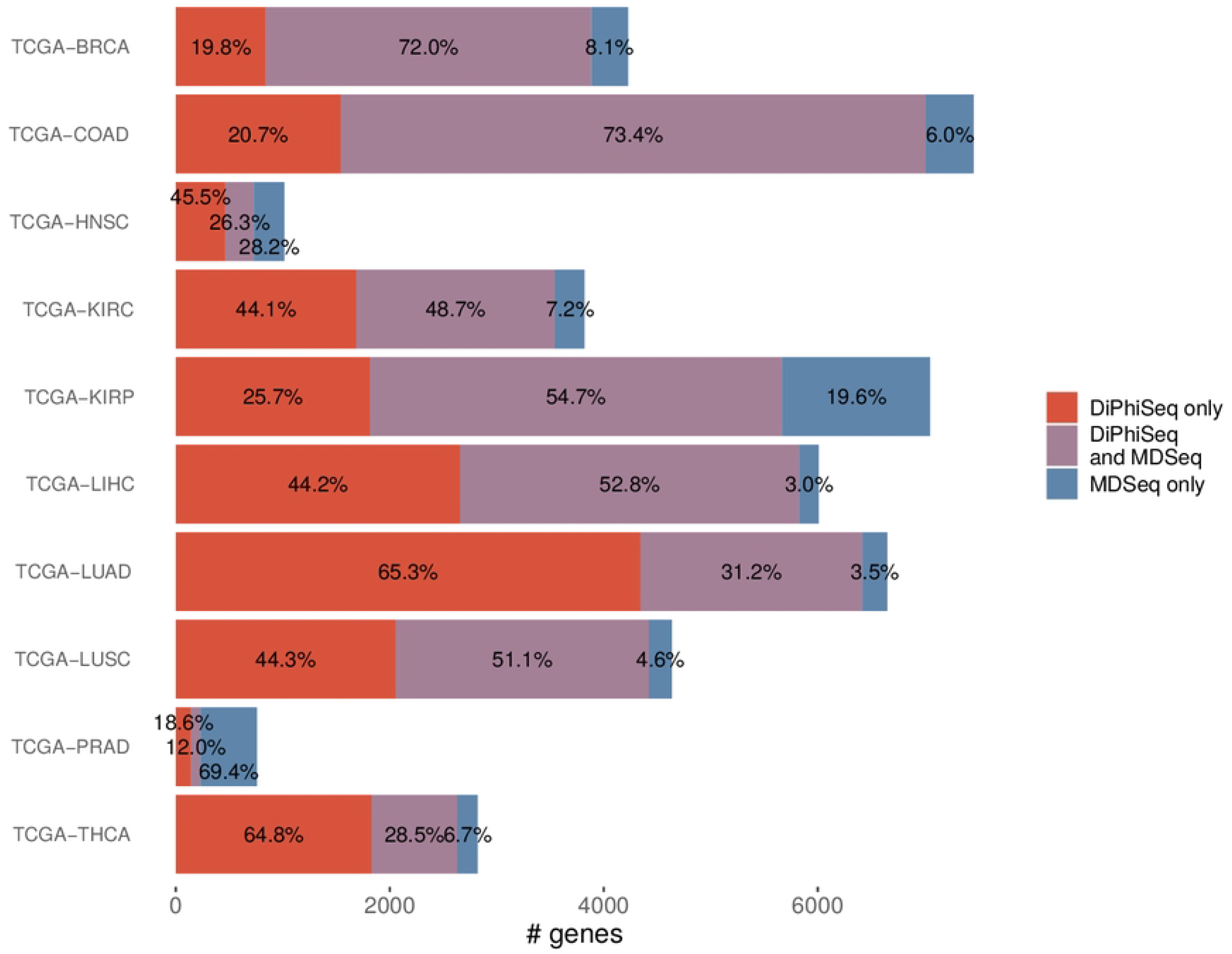
DD+ genes identified by either DiPhiSeq, MDSeq, or both among non-differentially expressed genes. Overdispersed genes in tumors (DD+) were identified by DiPhiSeq and MDSeq among non-differentially expressed genes for each TCGA dataset. Non-differentially expressed genes were identified by MDSeq.

For most of the analyzed datasets, the majority of the genes identified as DD+ are labeled as such by both methods (from 48.7% to 73.4%). DiPhiSeq identified most of the DD+ genes for the lung adenocarcinoma (TCGA-LUAD) and thyroid (TCGA-THCA) datasets (65.3% and 64.8%, respectively). Regarding the head and neck (TCGA-HNSC) and prostate (TCGA-PRAD) datasets, which are the two datasets for which only a few DD+ genes are detected, most of the DD+ genes are only identified with DiPhiSeq (45.5%) and MDSeq (69.4%), respectively. Overall, DiPhiSeq identifies more DD+ genes that are not detected by MDSeq than the other way around. This trend may be explained by the higher sensitivity of DiPhiSeq in detecting differential dispersion in expression data with large datasets, *i.e*. datasets composed of sets of at least 50 samples.

#### GO term enrichment analysis

To gain biological insight, an analysis of enrichment in Gene Ontology (GO) terms among DD+ genes for each TCGA dataset was conducted. To obtain the largest possible set of DD+ genes, we filtered highly DE genes with MDSeq using the maximum mean fold-change threshold values that maintain the FDR of the differential dispersion detection below 0.05 according to our simulation study, that is, 1.25 for the breast cancer dataset (TCGA-BRCA) and 1.30 for the others. Since these thresholds for the mean fold-change are quite low, we consider the genes that pass these filters to be lowly DE genes. Henceforth, DD+ genes refer to lowly DE, rather than non-DE, genes whose expression exhibits an increase of dispersion among tumor samples. The numbers of lowly DE genes and DD genes according to DiPhiSeq and MDSeq are displayed in S6 Fig, and the overlaps of the sets of DD+ genes are displayed in S7 Fig. Since DD+ genes are identified with an FDR below 0.05 with both methods according to our simulation study, the entire set of DD+ genes identified either by both methods or only one of them is taken into account to gain the most biological knowledge in the GO term enrichment analysis for each dataset. We used redundancy reduction methods to ease the comparison of enriched GO terms across all the analyzed datasets (see Methods for more details). The top 40 representative terms and the p-values of their enrichment in each dataset are shown in Figure 6. The full list of enriched representative GO terms is available in S1 File, and an overview is displayed in S8 Fig.

**Fig 6.**
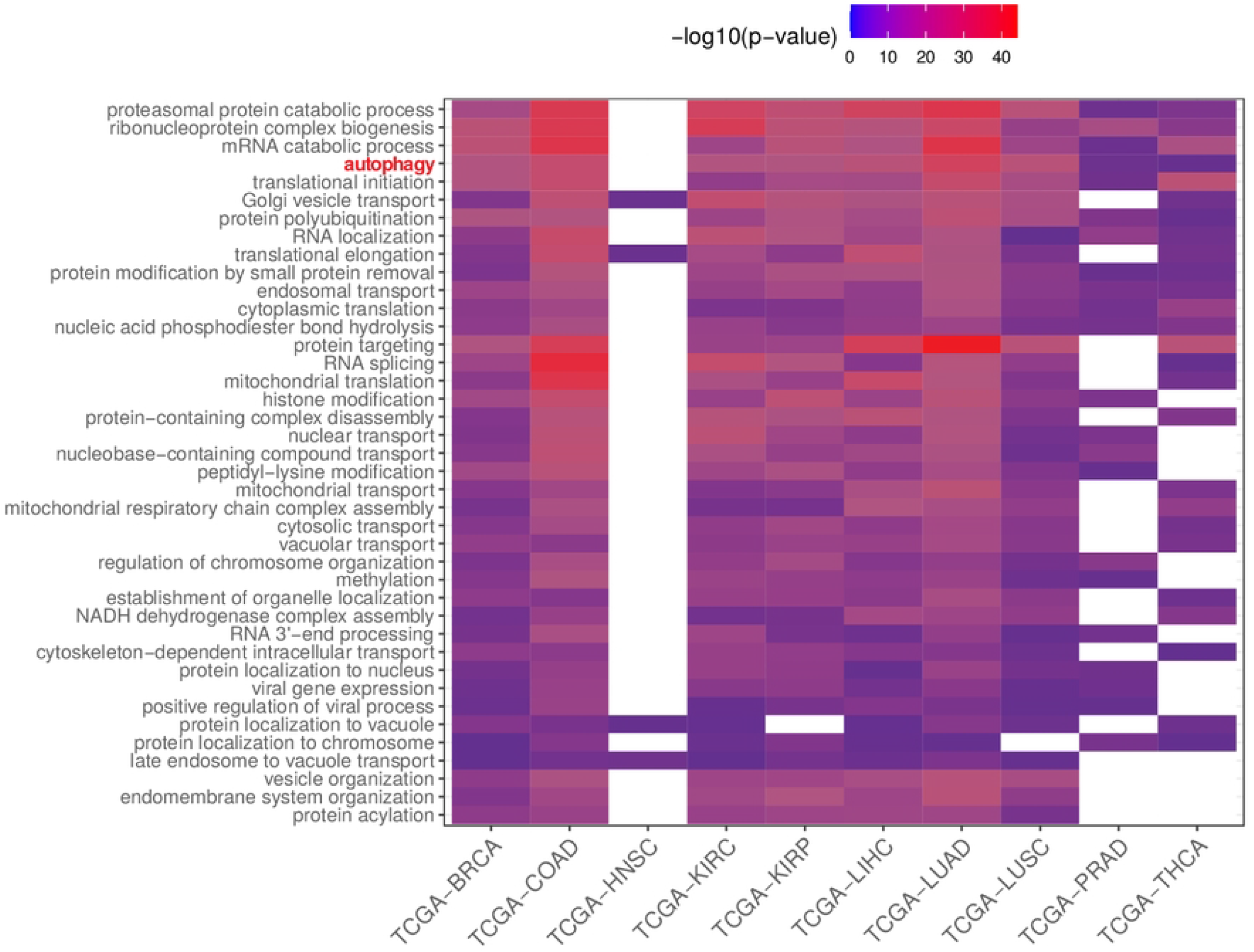
Enriched GO terms among DD+ genes according to DiPhiSeq and/or MDSeq for each TCGA dataset. Top 40 representative enriched Gene Ontology (GO) terms among overdispersed genes in tumors (DD+), ordered first by the number of datasets for which they are enriched (decreasing order) and second by the mean p-values of enrichment across all datasets (increasing order). Highly differentially expressed genes were filtered out using MDSeq, and DD+ genes were identified among lowly differentially expressed genes.

Interestingly, among DD+ genes, the most significantly enriched GO terms were the most widespread across all the analyzed tissues and focused on some key cellular functions, such as catabolism. In contrast, GO terms that were found to be significantly enriched for only a few datasets tended to have higher p-values than the most widely enriched GO terms (S8 Fig and S1 File). This striking result suggests some common features in tumoral development and progression, regardless of the tissue of origin, whose involved gene expression is characterized more by an increase in dispersion than by a change in the mean in tumors.

## Discussion

To our knowledge, our work is the first study to thoroughly assess the performance of methods to detect differential dispersion in RNA-seq data and, more generally, differential variance in gene expression data. We characterized DiPhiSeq and MDSeq performances based on simulated datasets. In particular, we identified key parameters to use to increase the sensitivity and to control the FDR. These simulations enabled us to propose recommendations to reliably apply these methods to real datasets.

### Gene expression dispersion in cancer

#### Overall increase of dispersion and robustness

By applying DiPhiSeq and MDSeq to TCGA datasets, we identified an overall increase in the dispersion in the expression of many lowly DE genes in tumors in comparison with normal tissues. Also analyzing TCGA datasets, Han *et al*. have already revealed an increase in the coefficient of variation of gene expression in tumors of breast, colon, lung and liver cancers [28]. In addition, using microarray data, Ho *et al*. [8] also noticed that an increase in gene expression variance in a disease condition such as cancer is more common than a decrease. Our work confirms these results, extends them to other cancers and increases their reliability by using RNA-seq data and methods based on a more appropriate statistical framework, and rigorously validates them in a simulation study. This increase in the dispersion in gene expression in tumors may reflect the huge variety of genetic perturbations occurring in their development and their polyclonal origin [48]. It may result from a loss of control of gene expression in cancer cells, *e.g*. loss of specificity in signaling cascades, transcriptional activity (cis and trans factors) or post-transcriptional regulation, *e.g*. splicing events or translation inhibition by microRNAs [49, 50]. Whatever its origin, this high variability in gene expression in cancer cells may be considered as a gain of robustness, as defined by Kitano [51]. The increase in the dispersion in the expression of hundreds of genes in tumors may enable them to adapt quickly and effectively to any perturbation of their environment. This may explain the resistance to treatment often observed, in particular to treatments that were effective during the first years of application [48]. These genes, whose mean expression does not vary significantly but whose dispersion of expression increases in cancer, form *de facto* a new space for the discovery of potential biomarkers.

#### Overrepresented functions among DD+ genes

We revealed that the biological processes that were the most significantly enriched among DD+ genes were also the most widespread across the different analyzed cancers. This striking result suggests common traits in tumoral development and progression pertaining to some key biological processes. It is worth noting that many of them are related to catabolism, *e.g*. “proteasomal protein catabolic process”, “mRNA catabolic process” or “protein targeting”, as previously shown by Han *et al*. [28]. In particular, several processes related to the ubiquitin-proteasome system, which is a major controller of the protein degradation process and is highly involved in cancer [52], are found among the most significant results (“protein polyubiquitination”, “proteasomal ubiquitin-independent protein catabolic process”). In contrast, no process related to anabolism was found among the most frequently enriched processes among DD+ genes, suggesting that catabolic processes are much more affected by the dysregulation of gene expression than are anabolic processes.

Autophagy was also found among the biological processes significantly enriched among DD+ genes for all the analyzed datasets. Similar to the proteasome, it is a main recycling system for biological molecules that enables cells to survive critical situations such as nutrient starvation and the degradation of damaged organelles or pathogens. In pre-malignant cells, autophagy actively acts to preserve the physiological homeostasis of multiple functions, *e.g*. elimination of mutagenic entities, decrease local inflammation, and thus aid the struggle against tumor development. In malignant cells, autophagy affects the tumor progession and the response to treatment in multiple ways, some of which act in opposition. Autophagy desensitizes cells to programmed cell death mediated by different treatment strategies but is also involved in danger signal emission which triggers an immune response through antigen presentation. Thus, the overall effect of autophagy on tumor progression and response to treatment is context-dependent [53]. The increase in the dispersion in the expression of genes involved in autophagic processes reveals the complexity of these processes in tumor progression and may lead one to wonder whether they should be induced or, on the contrary, inhibited as a cancer treatment [54]. Some treatments indeed aim to stimulate these processes, while others aim to inhibit them [55].

Our results revealed that the expression of the genes involved in these processes is mainly affected by an increase in dispersion in tumors rather than a change in the mean. Although DD genes are the main focus of interest in our approach, we also identified biological processes enriched among highly upregulated (S2 File) and highly downregulated genes (S3 File) in tumors with respect to healthy samples. Among all the previously discussed catabolic GO terms enriched among DD+ genes, the GO term “proteasomal ubiquitin-independent protein catabolic process” is the only one to also be enriched among highly upregulated genes (S2 File) for the single breast cancer dataset. Thus, these previously discussed catabolic GO terms are specifically enriched among DD+ genes for a large number of different tissues, which highlights the interest in searching for changes in dispersion, in addition to changes in the mean, to yield new insights into tumoral development and cancer treatment efficacy.

### Evaluation of DiPhiSeq and MDSeq differential dispersion detection performances

Based on our simulation study, we demonstrated that MDSeq must only be applied to lowly DE genes to reliably identify differential dispersion. In contrast, the detection of dispersion differences in gene expression data with DiPhiSeq is not affected by the presence of a difference in mean.

#### Dispersion and variance estimation

We showed that MDSeq tends to falsely identify differential dispersion among highly DE genes. The ability of MDSeq to predict differential dispersion for genes with opposite differential means is indeed poor, with a high level of false positives (S9 FigA). The GLM implemented in MDSeq is based on a reparameterization of the NB distribution, which has the advantage of explicitly modeling the variance of the random variable *Y*_*ig*_ describing the read counts but does not allow us to directly estimate the dispersion parameter of the classical NB distribution. Under this canonical model, the mean-variance relationship is defined by a quadratic function 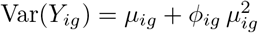. Thus, a change in variance may be due only to a change in mean, which explains why MDSeq achieves poor differential dispersion performance among highly DE genes but can still be reliably applied to identify differential dispersion among lowly DE genes, based on a nonsignificant p-value for the difference of mean test and a significant p-value for the difference of variance test (S9 FigB). In contrast, DiPhiSeq is based on the classical definition of the NB distribution and therefore allows us to estimate changes in dispersion. Our evaluations demonstrated that the detection of differential dispersion with DiPhiSeq is not sensitive to the presence of a mean fold-change and thus confirmed the claim of the DiPhiSeq authors: their methods effectively handle negatively associated mean and dispersion values [41]. Even if the main interest of searching for DD genes is to identify genes which are not detected by the classical differential expression analysis, *i.e*. non- or lowly DE genes, estimating differences in dispersion may help avoid misinterpreting a difference in dispersion as a difference in mean, and eventually bring new biological insights [56].

#### Specific features

One limitation of DiPhiSeq is that it does not allow the inclusion of any additional covariate in its statistical model to prevent some sources of bias from confounding the comparison of interest. This limitation is partly mitigated by the use of a Tukey’s biweight function that removes any aberrant value regardless of its source, either biological or technical. In contrast, similar to most differential expression analysis methods based on the NB distribution [37, 57], MDSeq implements a GLM that may take into account classical sources of bias, such as batch effects, in the detection of DD genes (see, for example, *LMAN2* expression in the lung adenocarcinoma dataset in S10 Fig) and therefore appears to better handle technical biases. Moreover, MDSeq also implements a zero-inflated model for which the goal is to control the statistical bias that may be introduced by an excess of null values in the analysis of gene expression data, which is particularly relevant for the analysis of single-cell expression data.

#### Computation time and large RNA-seq datasets

The computation time is another major difference between these two methods. The runtime of DiPhiSeq required to analyze the same datasets was indeed hundreds to thousand times longer than that of MDSeq (Table 2).

**Table 2.** DiPhiSeq and MDSeq computation times in minutes per dataset.

| Dataset | Sample sizes |  | Computation time (min) |  |
| --- | --- | --- | --- | --- |
|  |  |  | DiPhiSeq | MDSeq |
| <u>Simulated datasets:</u> |  |  |  |  |
|  | 30 | 30 | 339 | 2.3 |
|  | 40 | 40 | 451 | 2.6 |
|  | 50 | 50 | 563 | 3.2 |
|  | 100 | 100 | 1 140 | 5.5 |
| <u>TCGA datasets:</u> |  |  |  |  |
| TCGA-BRCA | 112 | 117 | 13 231 | 6.7 |
| TCGA-COAD | 41 | 46 | 4 135 | 4.1 |
| TCGA-HNSC | 43 | 43 | 6 372 | 3.5 |
| TCGA-KIRC | 72 | 72 | 9 766 | 4.7 |
| TCGA-KIRP | 31 | 31 | 4 422 | 3.3 |
| TCGA-LIHC | 50 | 50 | 6 192 | 3.4 |
| TCGA-LUAD | 57 | 67 | 5 427 | 4.9 |
| TCGA-LUSC | 49 | 49 | 7 417 | 3.9 |
| TCGA-PRAD | 52 | 54 | 6 528 | 3.7 |
| TCGA-THCA | 58 | 58 | 8 102 | 3.8 |

Computations were performed using a desktop computer with an Intel(R) Xeon(R) E3-1220 v5 @ 3.00 GHz 4-core CPU and 16 GB RAM. The first and second columns of the sample size part refer to samples from normal tissues and tumor tissues, respectively, for TCGA datasets. Mean computation times over replicates and all the evaluated maximum mean fold-change values for simulated datasets composed of lowly DE genes are displayed. MDSeq computation times for TCGA datasets pertain to mean and dispersion fold-changes above thresholds.

The possibility of using several CPUs and the use of optimization methods to estimate the GLM coefficients may explain, at least partly, the difference in computation time. Large gene expression datasets have become increasingly frequent with the development of comprehensive RNA-seq datasets, such as GTEx (Genotype-Tissue Expression) [58], and methods for integration of RNA-seq samples originating from heterogeneous sources while controlling technical biases [59]. The differential dispersion approach may therefore be applied in a variety of biological contexts. Thus, the computation time of DiPhiSeq may be a burden to its wide adoption.

### Gene expression variability at the single-cell level

Without any further specification, gene expression variability usually refers to cell-to-cell variability. Here, we analyzed RNA-seq data of samples composed of thousands of cells, *i.e*. bulk data coming from a population of different individuals. Some studies have demonstrated the limitations of inferences from bulk data regarding gene expression variability [60, 61]. Such approaches are unable to capture cell-to-cell variability and tend to average gene expression [62, 63]. Nevertheless, the estimation of gene expression based on this type of data may still exhibit some variability and provide a snapshot of the expression variability of a gene within populations of cells. We indeed identified a large number of genes with a significant change in dispersion in their expression between healthy and tumor bulk samples from different individuals. Single-cell RNA sequencing technologies have emerged in the last few years, enabling the study of gene expression variability at the cellular level. Their application in the differential dispersion approach is promising for a wide range of biological contexts. For example, in the context of cancer, the population of cells composing a tumor may exhibit a high level of gene expression variability, potentially leading to therapeutic failures [64]. However, analyzing the data generated with these techniques faces new methodological issues.

Technical null values, or dropouts, are much more present in single-cell RNA-seq data than in bulk RNA-seq data due to the limited amount of mRNA material available at the cellular level [65]. From the perspective of identifying DD genes using single-cell RNA-seq data, MDSeq seems to be currently more appropriate thanks to the development of dedicated features for managing excess technical null values or the incorporation of any technical bias through covariates in a GLM. Moreover, similar to comprehensive RNA-seq datasets, the high number of samples of single-cell RNA-seq datasets with respect to bulk RNA-seq datasets [65] may result in excessively long runtimes for DiPhiSeq, therefore hindering its application to such data.

## Conclusion

Overall, we showed that the application of the differential dispersion approach to gene expression data is relevant to gain knowledge of tumor progression and cancer treatment efficacy. With the emergence of comprehensive RNA-seq datasets, composed of either single-cell or bulk samples in a variety of biological contexts, we believe that the changes in dispersion in gene expression between samples from different conditions of biological interest should now be taken into account. In the classical differential mean expression analysis, it would provide a more realistic model of the data and, in the explicit goal of identifying genes with a differential variance in their expression, it may contribute to gaining new insights into biological processes and eventually to discovering new biomarkers and therapeutic avenues.

## Materials and methods

### RNA-seq dataset simulation

We simulated RNA-seq count datasets using the compcodeR R package [66]. The simulated datasets are composed of 10 000 genes and two sets of samples of equal size corresponding to biological conditions of interest. Read counts for gene *i* and sample *j* are generated by random sampling using a negative binomial distribution: *Y*_*ij*_ ∼ *𝒩 ℬ*(*μ*_*iρ*(*j*)_, *ϕ*_*iρ*(*j*)_), where *μ*_*iρ*(*j*)_ and *ϕ*_*iρ*(*j*)_ are the mean and dispersion parameters, respectively, and *ρ*(*j*) denotes the condition of sample *j* (*ρ*(*j*) ∈ {1; 2}). The mean *μ*_*i*1_ and dispersion *ϕ*_*i*1_ values for the first condition are estimated by pairs from real datasets [67, 68]. The mean and dispersion parameters for the second condition were generated by selecting a fraction of the genes to be subjected to a fold-change in mean or dispersion.

The dispersion of genes chosen to be differentially dispersed (DD) was determined as 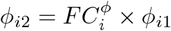, where 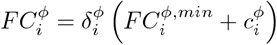, with 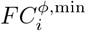 being a predefined minimum fold-change, 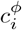 an extra amount drawn from an exponential distribution of parameter *λ* = 1, and 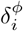 which is equally likely to be 1 or −1, so that half the genes have an increase in dispersion and the other half a decrease in dispersion in the second condition. We set 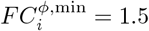 to ensure at least a 50% difference in dispersion between the two conditions. The non-DD genes have the same dispersion in both conditions: *ϕ*_*i*2_ = *ϕ*_*i*1_.

The mean expression of genes chosen to be differentially expressed (DE) was simulated according to two scenarios:

- Unconstrained mean expression fold-changes: Similar to what was done for the dispersion parameter of DD genes, 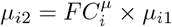, where 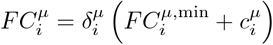. Several differences of mean minima were evaluated from 10% to 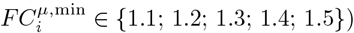, and 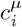 was drawn from an exponential distribution of parameter *λ* chosen in {0.85; 0.9; 1} depending on 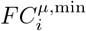 to obtain similar highest fold-change values (S1 Fig). Since the evaluation of differential mean expression detection performance is not the main goal of our approach, we simulated non-DE genes in a more realistic way than having the same mean expression value for both conditions. Instead, we allowed slight fold-changes by random sampling using uniform distributions: 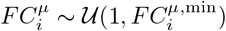, where the maximum value corresponds to the minimum value of fold-change for highly DE genes. These genes are therefore called lowly DE genes rather than non-DE genes.
- Moderated mean expression fold-changes: Since the purpose of these datasets is only to assess differential dispersion detection performance for lowly DE genes, the distinction between DE and non-DE genes is not required. For all the genes of these datasets, mean fold-changes were introduced using uniform distributions: 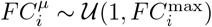, where 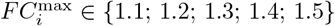.

For all the simulated datasets, the same fractions of DD genes (or non-DD genes) among highly DE genes and lowly DE genes (for the first set of simulations) were ensured, as well as the same fractions of DD genes with an increase in dispersion in the second condition (DD+) among upregulated genes (DE+) and downregulated genes (DE-) in the second condition. Thus, simulated datasets are composed of 50% DD genes and 50% non-DD genes and 50% highly DE genes and 50% lowly DE genes for the first set of simulations.

For the sake of realism, we introduced outliers with very high counts in all the simulated datasets since Li *et al*. [41] showed that they may have a dramatic effect on differential dispersion detection. Following the recommendations of Soneson and Delorenzi [44], we multiplied one read count by a value from 5 to 10 for 10% of the genes.

### RNA-seq data preprocessing

Before applying differential dispersion detection methods, classical RNA-seq data preprocessing steps were applied to all the simulated and TCGA datasets. First, read counts were normalized by the Trimmed Mean of M-values method [69, 70]. Lowly expressed genes were then independently filtered out using a threshold of 1 count per million [71].

### Differential dispersion detection

DiPhiSeq was applied to all the simulated and TCGA datasets with the default values for all the parameters, in particular the *c* parameter of Tukey’s biweight function set to 4 for both the mean and the dispersion estimation since the authors of DiPhiSeq found that this value enables robust parameter estimations [41]. The p-values for both differential mean and differential dispersion statistical tests were corrected by the Benjamini-Hochberg procedure [72] to control the FDR as recommended by the authors of DiPhiSeq.

The outlier removal function was applied with the minimum sample size lowered to 1 before applying the MDSeq main function to all the simulated and TCGA datasets. Batch effects were handled when analyzing TCGA datasets by supplying the sequencing runs that generated the RNA-seq samples, when available, as a covariate in the GLM for both the identification of DD genes and the filtering of highly DE genes. A range of threshold values from 1 to 1.5 was used to filter out highly DE genes, and DD genes were identified with a fold-change threshold of 1. The default values were used for the other parameters for both the outlier removal and MDSeq functions. The p-values for both differential mean and differential dispersion statistical tests were corrected by the Benjamini-Yekutieli procedure [73] to control the FDR as recommended by the authors of MDSeq.

We corrected the p-values using FDR-controlling procedures. As recommended by the authors of the respective methods, we corrected the p-values obtained with DiPhiSeq with the Benjamini-Hochberg procedure and those obtained with MDSeq using the Benjamini-Yekutieli procedure. We verified in both cases that this appropriately maintained the FDR of the differential dispersion detection below 0.05 (S2 Fig).

### Performance evaluation

The performances of differential expression and differential dispersion detection methods were evaluated based on the fold-changes of the mean and dispersion of expression introduced in the simulated datasets. The genes that were simulated to be highly DE or DD are the positive groups for the differential mean or the differential dispersion performance evaluations, respectively. The genes that were simulated to be lowly DE or non-DD are the corresponding negative groups. For both DiPhiSeq and MDSeq, a p-value for the differential mean or differential dispersion statistical test lower than 0.05 after the application of the appropriate FDR-controlling procedure was used to define positive detection. The comparisons with the positive groups enabled us to count true positive (TP) and false positive (FP) results for both differential expression and differential dispersion. Similarly, true negative (TN) and false negative (FN) results were identified thanks to a corrected p-value of differential mean or differential dispersion statistical test higher than 0.05 and the comparisons with the negative groups. The sensitivity (or true positive rate (TPR)), the false discovery rate (FDR) and the area under the ROC curve (AUC) were then computed based on these four categories of results.

### Gene Ontology term enrichment analysis across datasets

For each TCGA dataset, genes of interest, *e.g*. DD+ genes, were identified and Gene Ontology (GO) term enrichment analysis was performed using the Biological Processes (BP) ontology. Enriched GO terms were identified thanks to a hypergeometric test p-value after FDR control lower than 0.05 using the enrichGO function of the clusterProfiler R package [74]. The list of enriched GO terms was then reduced by gathering terms whose semantic similarity exceeded a threshold value. To do so, clusters of closely related GO terms were generated through the relevance method [75] to compute semantic similarity between GO terms. A high similarity threshold (0.8) was used to gather only closely related GO terms into clusters. The GO term whose p-value is the lowest among a cluster was then chosen to represent the entire cluster.

To facilitate comparisons across datasets, closely related GO terms were searched for among the previously simplified enriched GO term lists originating from each dataset. The similarity of all GO term pairs was calculated with the relevance method. These similarity values were then used to perform hierarchical clustering and gather closely related GO terms by using a conservative threshold value (0.8). For each cluster of closely related GO terms, the hierarchical structure of the BP ontology was used to identify a generic term common to all the GO terms. This common generic GO term was subsequently used as the representative term for the entire cluster, and its enrichment p-value was retrieved for each TCGA dataset containing an enriched GO term in the cluster.

## Supporting information

**S1 Fig. Fold-change distributions used to simulate differentially expressed genes**. Different minimum values *b* and extra amounts drawn from exponential distributions using different parameter values *λ* to have similar maximum values across all the distributions of fold-change values.

**S2 Fig. Correction of p-values for differential dispersion by the Benjamini-Hochberg and Benjamini-Yekutieli FDR-controlling procedures**. P-values obtained with DiPhiSeq (A) and MDSeq (B) for the detection of differential dispersion in gene expression data from a simulated dataset composed of two populations of 50 samples before and after correction by the Benjamini-Hochberg (BH) and Benjamini-Yekutieli (BY) procedures. All the genes are lowly DE with a mean fold-change of expression between 1 and 1.3. The red dotted lines represent a p-value threshold value of 0.05.

**S3 Fig. Differential dispersion performance with MDSeq after filtering highly DE genes using different fold-change threshold values**. (A) False discovery rate (FDR) and (B) true positive rate (TPR) values obtained with simulated datasets composed of two sample populations of equal size (panels on the horizontal axis). The maximum mean fold-change threshold values that enable an increase in the sensitivity of the differential dispersion detection while maintaining the FDR below 0.05 are displayed in red.

**S4 Fig. Differentially expressed and differentially dispersed genes according to DiPhiSeq and MDSeq for each TCGA dataset**. (A) Number of differentially expressed (DE) genes separated between those upregulated in tumors (DE+) and those downregulated in tumors (DE-), as detected by DiPhiSeq and MDSeq, per TCGA dataset. (B) Number of differentially dispersed (DD) genes separated between those overdispersed in tumors (DD+) and those underdispersed in tumors (DD-), as detected by DiPhiSeq and MDSeq, per TCGA dataset.

**S5 Fig. Differentially dispersed (DD) genes according to DiPhiSeq and MDSeq for each TCGA dataset**. (A) Number of overdispersed genes in tumors (DD+) and (B) number of underdispersed genes in tumors (DD-) separated between those upregulated in tumors (DE+), those downregulated in tumors (DE-) and those non-differentially expressed (non-DE), as detected by DiPhiSeq and MDSeq, per TCGA dataset.

**S6 Fig. Differentially dispersed genes among lowly differentially expressed genes for each TCGA dataset**. (A) Number of highly differentially expressed (DE) genes, separated between those upregulated in tumors (DE+) and those downregulated in tumors (DE-), and lowly DE genes, as detected by MDSeq, per TCGA dataset. Highly DE genes were filtered using the maximum mean fold-change threshold values, enabling us to keep the false discovery rate of the differential dispersion detection below 0.05 with respect to the number of available samples per dataset according to our simulation study. (B) Number of differentially dispersed (DD) genes among lowly DE genes separated between those overdispersed in tumors (DD+) and those underdispersed in tumors (DD-), as detected by DiPhiSeq and MDSeq, per TCGA dataset.

**S7 Fig. DD+ genes identified by either DiPhiSeq, MDSeq, or both among lowly differentially expressed genes**. Overdispersed genes in tumors (DD+) were identified by DiPhiSeq and MDSeq among the lowly differentially expressed (DE) genes for each TCGA dataset. Lowly DE genes were identified by MDSeq using the maximum mean fold-change threshold values, enabling us to keep the false discovery rate of the differential dispersion detection below 0.05 with respect to the number of available samples per dataset according to our simulation study.

**S8 Fig. Overview of the enriched GO terms among DD+ genes according to DiPhiSeq and/or MDSeq**. Overview of the entire list of representative enriched Gene Ontology (GO) terms among overdispersed genes in tumors (DD+), ordered first by the number of datasets for which they are enriched (decreasing order) and second by the mean p-values of enrichment across all datasets (increasing order). Highly differentially expressed genes were filtered out using MDSeq, and DD+ genes were identified among lowly differentially expressed genes.

**S9 Fig. Reliable identification of differentially dispersed genes among lowly differentially expressed genes with MDSeq**. Identification of differentially dispersed (DD) genes based on (A) a significant p-value for the difference of variance test or (B) a nonsignificant p-value for the difference of mean test and a significant p-value for the difference of variance test. True mean and dispersion log_2_-fold-changes (left panels) and estimated mean and variance log_2_-fold-changes with MDSeq (right panels) of a simulated dataset composed of two populations of 50 samples are displayed. Colours represent the results of the identification of differential dispersion by MDSeq using a log_2_-fold-change threshold of 0. The red dotted line is the *y* = *x* diagonal.

**S10 Fig. Batch effect handling by a covariate in the generalized linear model implemented by MDSeq**. Expression values of the *LMAN2* gene (lectin, mannose binding 2, ENSG00000169223) based on the TCGA dataset composed of samples from patients with lung adenocarcinoma (TCGA-LUAD) for whom both a tumor sample and a normal sample are available. Data are clustered according to sequencing batch. In batches 0946, 1107, 1206 and A277, which enabled the sequencing of only tumor samples, the dispersion of LMAN2 expression increased with respect to the other batches composed of samples from both conditions. Differential dispersion MDSeq p-values without the integration of batch effect by a blocking factor in the generalized linear model (GLM): 3.84 10^*−*4^; with the integration of batch effect by a blocking factor in the GLM: 1.36 10^*−*1^; DiPhiSeq differential dispersion p-value: 9.08 10^*−*5^.

**S1 File. Enriched GO terms among DD+ genes according to DiPhiSeq and/or MDSeq for each TCGA dataset**. List of representative enriched Gene Ontology (GO) terms among overdispersed genes in tumors (DD+), ordered first by the number of datasets for which they are enriched (decreasing order) and second by the mean p-values of enrichment across all datasets (increasing order). Highly differentially expressed genes were filtered out using MDSeq, and DD+ genes were identified among lowly differentially expressed genes.

**S2 File. Enriched GO terms among highly upregulated genes in tumors for each TCGA dataset**. List of representative enriched Gene Ontology (GO) terms among highly upregulated genes in tumors, ordered first by the number of datasets for which they are enriched (decreasing order) and second by the mean p-values of enrichment across all datasets (increasing order). Highly upregulated genes were identified by MDSeq using the maximum mean fold-change threshold values, enabling us to keep the FDR of the differential dispersion detection below 0.05, in agreement with our simulation study and with respect to the number of available samples per dataset, *i.e*. 1.25 for the breast cancer dataset (TCGA-BRCA) and 1.30 for the others.

**S3 File. Enriched GO terms among highly downregulated genes in tumors for each TCGA dataset**. List of representative enriched Gene Ontology (GO) terms among highly downregulated genes in tumors, ordered first by the number of datasets for which they are enriched (decreasing order) and second by the mean p-values of enrichment across all datasets (increasing order). Highly downregulated genes were identified by MDSeq using the maximum mean fold-change threshold values, enabling us to keep the FDR of the differential dispersion detection below 0.05, in agreement with our simulation study and with respect to the number of available samples per dataset, *i.e*. 1.25 for the breast cancer dataset (TCGA-BRCA) and 1.30 for the others.

## Acknowledgments

We thank Adeline Leclercq-Samson and Christophe Battail for their helpful recommendations.

## Notes

### Competing Interest Statement

The authors have declared no competing interest.

## References

1. Shendure J, Lieberman Aiden E. The expanding scope of DNA sequencing. Nat Biotechnol. 2012;30(11):1084–1094.

2. Peixoto A, Monteiro M, Rocha B, Veiga-Fernandes H. Quantification of multiple gene expression in individual cells. Genome Res. 2004;14(10A):1938–1947.

3. Son CG, Bilke S, Davis S, Greer BT, Wei JS, Whiteford CC, et al. Database of mRNA gene expression profiles of multiple human organs. Genome Res. 2005;15(3):443–450.

4. Escrich E, Moral R, García G, Costa I, Sánchez JA, Solanas M. Identification of novel differentially expressed genes by the effect of a high-fat n-6 diet in experimental breast cancer. Mol Carcinog. 2004;40(2):73–78.

5. Shen Y, Wang X, Jin Y, Lu J, Qiu G, Wen X. Differentially expressed genes and interacting pathways in bladder cancer revealed by bioinformatic analysis. Mol Med Rep. 2014;10(4):1746–1752.

6. Altintas DM, Allioli N, Decaussin M, de Bernard S, Ruffion A, Samarut J, et al. Differentially expressed androgen-regulated genes in androgen-sensitive tissues reveal potential biomarkers of early prostate cancer. PLoS One. 2013;8(6):e66278.

7. Emmert-Streib F, Tripathi S, de Matos Simoes R. Harnessing the complexity of gene expression data from cancer: from single gene to structural pathway methods. Biol Direct. 2012;7:44.

8. Ho JW, Stefani M, dos Remedios CG, Charleston MA. Differential variability analysis of gene expression and its application to human diseases. Bioinformatics. 2008;24(13):i390–398.

9. Eisenberg E, Levanon EY. Human housekeeping genes, revisited. Trends Genet. 2013;29(10):569–574.

10. Alemu EY, Carl JW, Corrada Bravo H, Hannenhalli S. Determinants of expression variability. Nucleic Acids Res. 2014;42(6):3503–3514.

11. Zhang M, Yao C, Guo Z, Zou J, Zhang L, Xiao H, et al. Apparently low reproducibility of true differential expression discoveries in microarray studies. Bioinformatics. 2008;24(18):2057–2063.

12. Suter DM, Molina N, Gatfield D, Schneider K, Schibler U, Naef F. Mammalian genes are transcribed with widely different bursting kinetics. Science. 2011;332(6028):472–474.

13. Losick R, Desplan C. Stochasticity and cell fate. Science. 2008;320(5872):65–68.

14. Cağatay T, Turcotte M, Elowitz MB, Garcia-Ojalvo J, Süel GM. Architecture-dependent noise discriminates functionally analogous differentiation circuits. Cell. 2009;139(3):512–522.

15. Viñuelas J, Kaneko G, Coulon A, Vallin E, Morin V, Mejia-Pous C, et al. Quantifying the contribution of chromatin dynamics to stochastic gene expression reveals long, locus-dependent periods between transcriptional bursts. BMC Biol. 2013;11:15.

16. Chalancon G, Ravarani CN, Balaji S, Martinez-Arias A, Aravind L, Jothi R, et al. Interplay between gene expression noise and regulatory network architecture. Trends Genet. 2012;28(5):221–232.

17. Volfson D, Marciniak J, Blake WJ, Ostroff N, Tsimring LS, Hasty J. Origins of extrinsic variability in eukaryotic gene expression. Nature. 2006;439(7078):861–864.

18. Pedraza JM, van Oudenaarden A. Noise propagation in gene networks. Science. 2005;307(5717):1965–1969.

19. Gerlinger M, Rowan AJ, Horswell S, Math M, Larkin J, Endesfelder D, et al. Intratumor heterogeneity and branched evolution revealed by multiregion sequencing. N Engl J Med. 2012;366(10):883–892.

20. Almendro V, Cheng YK, Randles A, Itzkovitz S, Marusyk A, Ametller E, et al. Inference of tumor evolution during chemotherapy by computational modeling and in situ analysis of genetic and phenotypic cellular diversity. Cell Rep. 2014;6(3):514–527.

21. Marusyk A, Almendro V, Polyak K. Intra-tumour heterogeneity: a looking glass for cancer? Nat Rev Cancer. 2012;12(5):323–334.

22. Wang K, Vijay V, Fuscoe JC. Stably Expressed Genes Involved in Basic Cellular Functions. PLoS One. 2017;12(1):e0170813.

23. Hasegawa Y, Taylor D, Ovchinnikov DA, Wolvetang EJ, de Torrenté L, Mar JC. Variability of Gene Expression Identifies Transcriptional Regulators of Early Human Embryonic Development. PLoS Genet. 2015;11(8):e1005428.

24. Wang K, Phillips CA, Rogers GL, Barrenas F, Benson M, Langston MA. Differential Shannon entropy and differential coefficient of variation: alternatives and augmentations to differential expression in the search for disease-related genes. Int J Comput Biol Drug Des. 2014;7(2-3):183–194.

25. Ecker S, Pancaldi V, Rico D, Valencia A. Higher gene expression variability in the more aggressive subtype of chronic lymphocytic leukemia. Genome Med. 2015;7(1):8.

26. Zhang F, Yao Shugart Y, Yue W, Cheng Z, Wang G, Zhou Z, et al. Increased Variability of Genomic Transcription in Schizophrenia. Sci Rep. 2015;5:17995.

27. Mason EA, Mar JC, Laslett AL, Pera MF, Quackenbush J, Wolvetang E, et al. Gene expression variability as a unifying element of the pluripotency network. Stem Cell Reports. 2014;3(2):365–377.

28. Han R, Huang G, Wang Y, Xu Y, Hu Y, Jiang W, et al. Increased gene expression noise in human cancers is correlated with low p53 and immune activities as well as late stage cancer. Oncotarget. 2016;7(44):72011–72020.

29. Mar JC, Matigian NA, Mackay-Sim A, Mellick GD, Sue CM, Silburn PA, et al. Variance of gene expression identifies altered network constraints in neurological disease. PLoS Genet. 2011;7(8):e1002207.

30. Zwiener I, Frisch B, Binder H. Transforming RNA-Seq data to improve the performance of prognostic gene signatures. PLoS One. 2014;9(1):e85150.

31. Marioni JC, Mason CE, Mane SM, Stephens M, Gilad Y. RNA-seq: an assessment of technical reproducibility and comparison with gene expression arrays. Genome Res. 2008;18(9):1509–1517.

32. O’Hara RB, Kotze DJ. Do not log-transform count data. Methods in Ecology and Evolution. 2010;1(2):118–122.

33. Conesa A, Madrigal P, Tarazona S, Gomez-Cabrero D, Cervera A, McPherson A, et al. A survey of best practices for RNA-seq data analysis. Genome Biol. 2016;17:13.

34. Robinson MD, Smyth GK. Moderated statistical tests for assessing differences in tag abundance. Bioinformatics. 2007;23(21):2881–2887.

35. Robinson MD, Smyth GK. Small-sample estimation of negative binomial dispersion, with applications to SAGE data. Biostatistics. 2008;9(2):321–332.

36. McCarthy DJ, Chen Y, Smyth GK. Differential expression analysis of multifactor RNA-Seq experiments with respect to biological variation. Nucleic Acids Res. 2012;40(10):4288–4297.

37. Love MI, Huber W, Anders S. Moderated estimation of fold change and dispersion for RNA-seq data with DESeq2. Genome Biol. 2014;15(12):550.

38. Wu H, Wang C, Wu Z. A new shrinkage estimator for dispersion improves differential expression detection in RNA-seq data. Biostatistics. 2013;14(2):232–243.

39. Landau WM, Liu P. Dispersion estimation and its effect on test performance in RNA-seq data analysis: a simulation-based comparison of methods. PLoS One. 2013;8(12):e81415.

40. Ran D, Daye ZJ. Gene expression variability and the analysis of large-scale RNA-seq studies with the MDSeq. Nucleic Acids Res. 2017;45(13):e127.

41. Li J, Lamere AT. DiPhiSeq: robust comparison of expression levels on RNA-Seq data with large sample sizes. Bioinformatics. 2019;35(13):2235–2242.

42. Yu L, Fernandez S, Brock G. Power analysis for RNA-Seq differential expression studies. BMC Bioinformatics. 2017;18(1):234.

43. Bonafede E, Picard F, Robin S, Viroli C. Modeling overdispersion heterogeneity in differential expression analysis using mixtures. Biometrics. 2016;72(3):804–814.

44. Soneson C, Delorenzi M. A comparison of methods for differential expression analysis of RNA-seq data. BMC Bioinformatics. 2013;14:91.

45. Bi R, Liu P. Sample size calculation while controlling false discovery rate for differential expression analysis with RNA-sequencing experiments. BMC Bioinformatics. 2016;17:146.

46. Hwang YT, Chu SK, Ou ST. Evaluations of FDR-controlling procedures in multiple hypothesis testing. Statistics and Computing. 2011;21(4):569–583.

47. Institute NC. The Cancer Genome Atlas; 2017. https://cancergenome.nih.gov/.

48. Burrell RA, Swanton C. Tumour heterogeneity and the evolution of polyclonal drug resistance. Mol Oncol. 2014;8(6):1095–1111.

49. Schmiedel JM, Klemm SL, Zheng Y, Sahay A, Blüthgen N, Marks DS, et al. Gene expression. MicroRNA control of protein expression noise. Science. 2015;348(6230):128–132.

50. Yang Z, Dong D, Zhang Z, Crabbe MJ, Wang L, Zhong Y. Preferential regulation of stably expressed genes in the human genome suggests a widespread expression buffering role of microRNAs. BMC Genomics. 2012;13 Suppl 7:S14.

51. Kitano H. Biological robustness. Nat Rev Genet. 2004;5(11):826–837.

52. Shen M, Schmitt S, Buac D, Dou QP. Targeting the ubiquitin-proteasome system for cancer therapy. Expert Opin Ther Targets. 2013;17(9):1091–1108.

53. Rybstein MD, Bravo-San Pedro JM, Kroemer G, Galluzzi L. The autophagic network and cancer. Nat Cell Biol. 2018;20(3):243–251.

54. Poillet-Perez L, Sarry JE, Joffre C. Autophagy is a major metabolic regulator involved in cancer therapy resistance. Cell Rep. 2021;36(7):109528.

55. Levy JMM, Towers CG, Thorburn A. Targeting autophagy in cancer. Nat Rev Cancer. 2017;17(9):528–542.

56. Cui W, Xue H, Wei L, Jin J, Tian X, Wang Q. High heterogeneity undermines generalization of differential expression results in RNA-Seq analysis. Hum Genomics. 2021;15(1):7.

57. Robinson MD, McCarthy DJ, Smyth GK. edgeR: a Bioconductor package for differential expression analysis of digital gene expression data. Bioinformatics. 2010;26(1):139–140.

58. Battle A, Brown CD, Engelhardt BE, Montgomery SB, Aguet F, Ardlie KG, et al. Genetic effects on gene expression across human tissues. Nature. 2017;550(7675):204–213.

59. Wang Q, Armenia J, Zhang C, Penson AV, Reznik E, Zhang L, et al. Unifying cancer and normal RNA sequencing data from different sources. Sci Data. 2018;5:180061.

60. Bengtsson M, Ståhlberg A, Rorsman P, Kubista M. Gene expression profiling in single cells from the pancreatic islets of Langerhans reveals lognormal distribution of mRNA levels. Genome Res. 2005;15(10):1388–1392.

61. Mar JC, Rubio R, Quackenbush J. Inferring steady state single-cell gene expression distributions from analysis of mesoscopic samples. Genome Biol. 2006;7(12):R119.

62. Piras V, Selvarajoo K. The reduction of gene expression variability from single cells to populations follows simple statistical laws. Genomics. 2015;105(3):137–144.

63. Richard A, Boullu L, Herbach U, Bonnafoux A, Morin V, Vallin E, et al. Single-Cell-Based Analysis Highlights a Surge in Cell-to-Cell Molecular Variability Preceding Irreversible Commitment in a Differentiation Process. PLoS Biol. 2016;14(12):e1002585.

64. Horning AM, Wang Y, Lin CK, Louie AD, Jadhav RR, Hung CN, et al. Single-Cell RNA-seq Reveals a Subpopulation of Prostate Cancer Cells with Enhanced Cell-Cycle-Related Transcription and Attenuated Androgen Response. Cancer Res. 2018;78(4):853–864.

65. Haque A, Engel J, Teichmann SA, Lönnberg T. A practical guide to single-cell RNA-sequencing for biomedical research and clinical applications. Genome Med. 2017;9(1):75.

66. Soneson C. compcodeR–an R package for benchmarking differential expression methods for RNA-seq data. Bioinformatics. 2014;30(17):2517–2518.

67. Pickrell JK, Marioni JC, Pai AA, Degner JF, Engelhardt BE, Nkadori E, et al. Understanding mechanisms underlying human gene expression variation with RNA sequencing. Nature. 2010;464(7289):768–772.

68. Cheung VG, Nayak RR, Wang IX, Elwyn S, Cousins SM, Morley M, et al. Polymorphic cis- and trans-regulation of human gene expression. PLoS Biol. 2010;8(9).

69. Robinson MD, Oshlack A. A scaling normalization method for differential expression analysis of RNA-seq data. Genome Biol. 2010;11(3):R25.

70. Dillies MA, Rau A, Aubert J, Hennequet-Antier C, Jeanmougin M, Servant N, et al. A comprehensive evaluation of normalization methods for Illumina high-throughput RNA sequencing data analysis. Brief Bioinform. 2013;14(6):671–683.

71. Bourgon R, Gentleman R, Huber W. Independent filtering increases detection power for high-throughput experiments. Proc Natl Acad Sci U S A. 2010;107(21):9546–9551.

72. Benjamini Y, Hochberg Y. Controlling the False Discovery Rate: A Practical and Powerful Approach to Multiple Testing. Journal of the Royal Statistical Society Series B (Methodological). 1995;57(1):289–300.

73. Benjamini Y, Yekutieli D. The control of the false discovery rate in multiple testing under dependency. Ann Statist. 2001;29(4):1165–1188.

74. Yu G, Wang LG, Han Y, He QY. clusterProfiler: an R package for comparing biological themes among gene clusters. OMICS. 2012;16(5):284–287.

75. Schlicker A, Domingues FS, Rahnenführer J, Lengauer T. A new measure for functional similarity of gene products based on Gene Ontology. BMC Bioinformatics. 2006;7:302.

